# Molecular mechanism of evolution and human infection with the novel coronavirus (2019-nCoV)

**DOI:** 10.1101/2020.02.17.952903

**Authors:** Jiahua He, Huanyu Tao, Yumeng Yan, Sheng-You Huang, Yi Xiao

## Abstract

Since December, 2019, an outbreak of pneumonia caused by the new coronavirus (2019-nCoV) has hit the city of Wuhan in the Hubei Province. With the continuous development of the epidemic, it has become a national public health crisis and calls for urgent antiviral treatments or vaccines. The spike protein on the coronavirus envelope is critical for host cell infection and virus vitality. Previous studies showed that 2019-nCoV is highly homologous to human SARS-CoV and attaches host cells though the binding of the spike receptor binding domain (RBD) domain to the angiotensin-converting enzyme II (ACE2). However, the molecular mechanisms of 2019-nCoV binding to human ACE2 and evolution of 2019-nCoV remain unclear. In this study, we have extensively studied the RBD-ACE2 complex, spike protein, and free RBD systems of 2019-nCoV and SARS-CoV using protein-protein docking and molecular dynamics (MD) simulations. It was shown that the RBD-ACE2 binding free energy for 2019-nCoV is significantly lower than that for SARS-CoV, which is consistent with the fact that 2019-nCoV is much more infectious than SARS-CoV. In addition, the spike protein of 2019-nCoV shows a significantly lower free energy than that of SARS-CoV, suggesting that 2019-nCoV is more stable and able to survive a higher temperature than SARS-CoV. This may also provide insights into the evolution of 2019-nCoV because SARS-like coronaviruses are thought to have originated in bats that are known to have a higher body-temperature than humans. It was also revealed that the RBD of 2019-nCoV is much more flexible especially near the binding site and thus will have a higher entropy penalty upon binding ACE2, compared to the RBD of SARS-CoV. That means that 2019-nCoV will be much more temperature-sensitive in terms of human infection than SARS-CoV. With the rising temperature, 2019-nCoV is expected to decrease its infection ability much faster than SARS-CoV, and get controlled more easily. The present findings are expected to be helpful for the disease prevention and control as well as drug and vaccine development of 2019-nCoV.

## 1 INTRODUCTION

Coronaviruses (CoVs) are a group of enveloped positive-stranded RNA viruses that can cause respiratory, intestinal and central nervous system infections in humans and animals [1]. Until last year, six strains of coronaviruses that are able to infect humans have been identified [1, 2]. Among them, four human coronaviruses, including HCoV-OC43, HCoV-229E, HCoV-NL63, and HCoVHKU1, are not highly pathogenic and only cause mild respiratory diseases [1]. However, two other coronaviruses, the severe acute respiratory syndrome coronavirus (SARS-CoV) [3–6] and the Middle East respiratory syndrome coronavirus (MERS-CoV) [7, 8], have caused two large-scale pandemics and resulted in more than 8000 cases including nearly 800 related deaths and about 2500 cases including about 860 related deaths, respectively. The outbreaks of SARS-CoV and MERS-CoV showed that some coronaviruses can be highly-pathogenic viruses when they transmit to humans from animals [9]. Therefore, it is urgent to develop antiviral treatments or vaccines targeting such high-risk coronaviruses like SARS-CoV and MERS-CoV.

Before efficient antiviral drugs or vaccines are developed for SARS-CoV or MERS-CoV, another outbreak of pneumonia caused by a new coronavirus (2019-nCoV) has emerged in Wuhan since December 2019. As of February 17, 2020, 2019-nCoV has caused more than 70000 cases in China and nearly 8000 cases around the world with about 1700 related deaths [10–14]. The full-length genome sequence of 2019-nCoV was soon determined by the Zhang group [15]. It was revealed that 2019-nCoV has a probable bat origin and is 96% identical at the whole-genome level to a bat SARS-like coronavirus [16]. In addition, 2019-nCoV is also closely related to other SARS-like coronaviruses and shares 79.5% sequence identify to SARS-CoV [16]. For some encoded proteins like coronavirus main proteinase (3CLpro), papain-like protease (PLpro), and RNA-dependent RNA polymerase (RdRp), the sequence identity is even higher and can be as high as 96% between 2019-nCoV and SARS-CoV [17]. Therefore, it has been thought that 2019-nCoV would function the similar way to SARS-CoV in the human-infection and pathogenic mechanism [16–18].

Coronaviruses uses the surface spike (S) glycoprotein on the envelope to attach host cells and mediate host cell membrane and viral membrane fusion during infection [19]. The spike protein includes two regions, S1 and S2, where S1 is for host cell receptor binding and S2 is for membrane fusion. The S1 region also includes an N-terminal domain (NTD) and three C-terminal domains (CTD1, CTD2, and CTD3) [21]. For SARS-CoV, the receptor binding domain (RBD) is located in the CTD1 of the S1 region. SARS-CoV attaches the human host cells through the binding of the RBD protein to the angiotensin-converting enzyme II (ACE2) [20]. Therefore, the interaction between RBD and ACE2 is a prerequisite for the human infection with SARS-CoV. Given the high homology between SARS-CoV and 2019-nCoV, it was expected that 2019-nCoV would also use the ACE2 molecule as the receptor to entry human cells [18]. This hypothesis was further experimentally confirmed by the virus infectivity studies from the Shi group, in which 2019-nCoV is able to use the ACE2 proteins from humans, Chinese horseshoe bats, and civet as an entry receptor in the ACE2-expressing cells, but not cells without ACE2 [16]. Xu et al. have used MOE to calculate the binding free energies between the RBD of spike protein and human ACE2, showing that the binding free energy between 2019-nCoV and human ACE2 was −50.6 kcal/mol, whereas that between SARS-CoV and human ACE2 was −78.6 kcal/mol [18]. Very recently, the Cryo-EM structure of the 2019-nCoV spike protein in the prefusion conformation has been determined [22]. The biophysical and structural evidence suggested that 2019-nCoV binds ACE2 with higher affinity than SARS-CoV [22].

Although it seems clear that 2019-nCoV infects human cells through the binding of the RBD domain to the human ACE2 receptor [16–18, 22], the molecular mechanism of the binding between the RBD protein and the ACE2 receptor is still unknown. Many questions remain to be answered. For example, previous studies showed that the binding affinity between 2019-nCoV and human ACE2 is weaker than that between SARS-CoV and human ACE2 [18]. However, in reality, 2019-nCoV has resulted in many more cases and seems to be more infectious than SARS-CoV. In this study, we have extensively investigated the spike protein/human ACE2 protein systems of 2019-nCoV and SARS-CoV by using protein-protein docking and molecular dynamics (MD) simulations. Specifically, we have extensively studied the free energies and dynamics of RBD-ACE2 binding, spike protein, and free RBD systems. Given that the spike protein is not only a potential drug target but also the virus antigen, the present study will be beneficial for the drug design, vaccine development, and disease prevention for 2019-nCoV.

## 2 MATERIALS AND METHODS

### 2.1 Structure preparation

In this study, we have investigated the RBD-ACE2 complex, spike protein, and free RBD systems of SARS-CoV (GenBank ID: NP 828851.1) and 2019-nCoV (GenBank ID: MN908947.3). For SARS-CoV, the RBD-ACE2 complex structure was directly downloaded from the Protein Data Bank (PDB entry: 3SCI) [23]. Then, all the water molecules were removed from the complex structure. The free RBD structure was obtained by removing the ACE2 protein from the RBD-ACE2 complex. The structure of the trimeric spike protein of SARS-CoV was also downloaded from the PDB (PDB entry: 6ACD) [21]. For 2019-nCoV, the three dimensional (3D) RBD structure was modeled based on the RBD structure of SARS-CoV using the MODELLER program [24], where the sequence alignment was performed using the ClustalW program [25, 26]. The complex structure between the 2019-nCoV RBD protein and human ACE2 was then predicted by our protein-protein docking approach [28–30]. The 3D structure of the trimeric spike protein for 2019-nCoV was constructed based on the structure of the SARS-CoV spike protein using MODELLER.

### 2.2 Protein-protein docking

The complex structure between the 2019-nCoV RBD protein and the human ACE2 molecule was predicted using our hybrid protein-protein docking algorithm, HDOCK [27–30]. Specifically, given the individual structures of the spike RBD protein and the human ACE2 molecule, HDOCK will globally sample all possible binding modes between the two proteins through a fast Fourier transform (FFT) search strategy [30]. Then, all the sampled binding modes were evaluated by our iterative knowledge-based scoring function [31]. Last, the binding modes were ranked according to their binding energy scores, and the top ten binding modes were provided to users. During the docking calculation, all the default parameters were used. Namely, the grid spacing was set to 1.2 Å for 3D translational search, the angle interval was set to 15° for rotational sampling in 3D Euler space, the binding interface information in the PDB was automatically applied during the modeling of individual structures. A web server version of our HDOCK algorithm can be freely accessed from our web site at http://hdock.phys.hust.edu.cn/ [29].

### 2.3 MD simulations

The AMBER suite was used for the MD simulations [32]. Before the simulations, the missing residues in the middle of a chain were added using the MODELLER [24]. During the simulations, the ff14SB force field was selected [33], explicit solvent model was used, the time step was set to 2 fs, langevin dynamics were used for temperature control, and the program “pmemd.cuda” was used as the simulation engine, where the simulations were performed on a GPU compute node [34]. Specifically, for each system, the following four stages of MD simulations were performed before the production simulation: (1) A 1000-step simulation was first run to minimize the solvated protein system with weakly restraints on the backbone atoms; (2) The system was then heated to 300K by a 25000-step (i.e. 50ps) simulation with weakly restraints on the backbone atoms; (3) Next, another 25000-step (i.e. 50ps) constant pressure simulation was conducted to equilibrating the density of the system at 300 K; (4) The system was then equilibrated by a 250000-step (i.e. 500ps) of constant pressure simulation at 300K. Finally, two 2500000-step production simulations were run to record the trajectories of the system at 300K, where the coordinates were written out every 5000 steps (i.e. 10ps), resulting in a total of 10ns simulation with 1000 recorded trajectories. After the simulations, the “MMPBSA.py” was used to calculate free energies of the systems using the MM-GBSA model [35], and the “cpptraj” was used to analyze the coordinate trajectories [36].

## 3 RESULTS AND DISCUSSION

### 3.1 The RBD-ACE2 docking

The RBD proteins of 2019-nCoV and SARS-CoV exhibit a high sequence similarity (89.2%) with a sequence identity of 73.7%. The high homology resulted in an accurate 3D RBD model of 2019-nCoV with a small RMSD of 0.55 Å from the experimental SARS-CoV RBD structure. With the experimental human ACE2 structure and the 2019-nCoV RBD model, we then performed protein-protein docking to predict their binding mode using our HDOCK approach [28–30]. Figure 1A shows the predicted complex structure between the human ACE2 molecule and the 2019-nCoV RBD protein. It can be seen from the figure that the predicted RBD-ACE2 complex structure for 2019-nCoV is very close to the experimentally determined RBD-ACE2 complex structure for SARS-CoV, and the interface root-mean-square-deviation (RMSD) between the two complexes is 0.473 Å, showing that the RBDs of 2019-nCoV and SARS-CoV bind to the same site of the human ACE2 receptor (Figure 1A). These results can also be understood by comparing the residues at the RBD-ACE2 binding interface for 2019-nCoV and SARS-CoV. The binding sites on the RBD proteins of 2019-nCoV and SARS-CoV are very conserved and the corresponding residues show a high sequence similarity of 83.3% (Figures 1B and C). Among them, those hydrophobic residues that are important for protein-protein interactions are especially conserved. For the RBD of 2019-nCoV, there are 13 hydrophobic residues at the binding site, which are comparable to 13 hydrophobic residues for that of SARS-CoV (Figure 1B).

**Figure 1:**
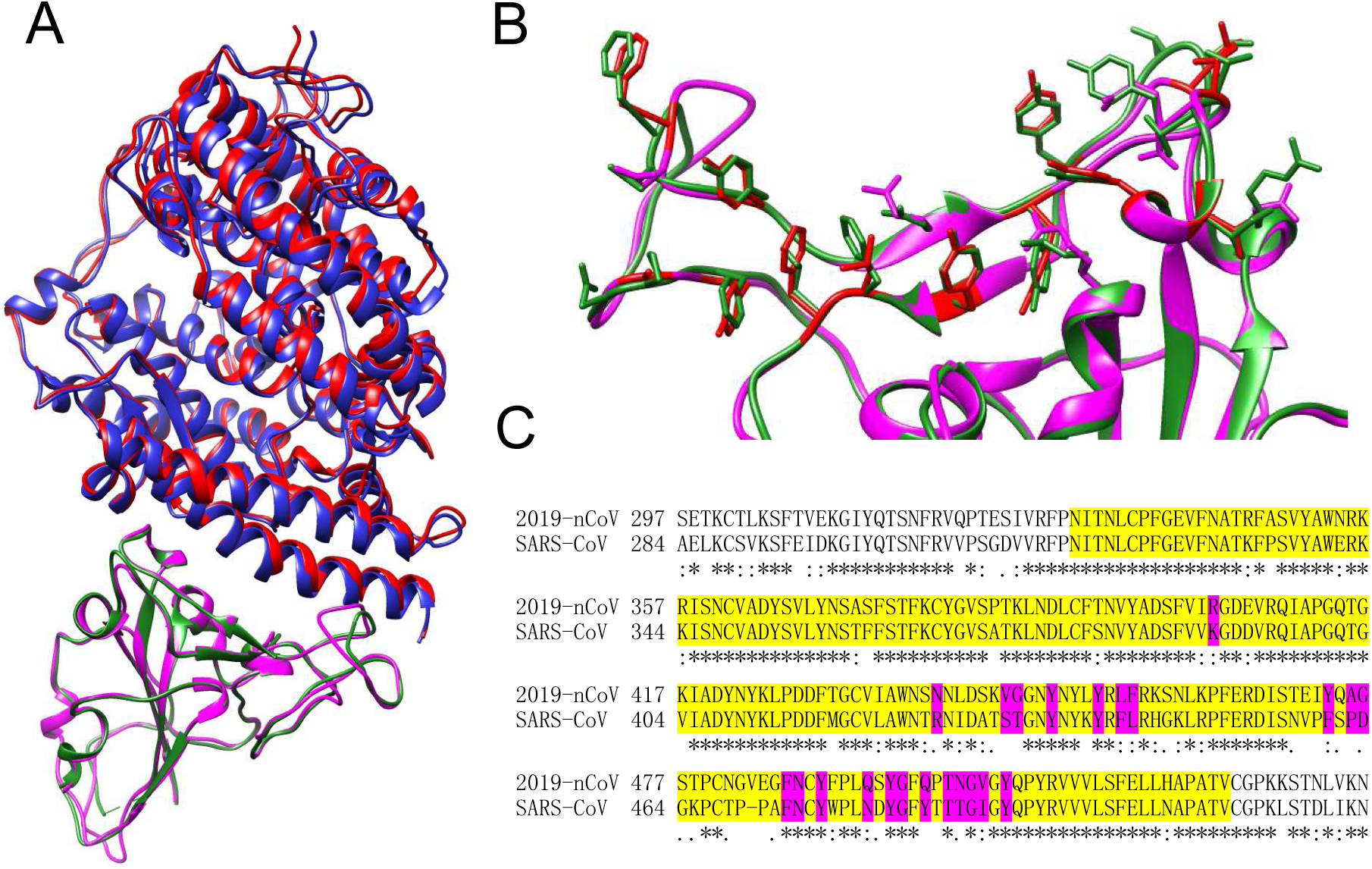
Protein-protein docking with the 2019-nCoV RBD and human ACE2. **(A)** The comparison between the predicted 2019-nCoV RBD-ACE2 complex and the experimental SARS-CoV RBD-ACE2 structure (3SCI:A/E), where the RBD and ACE2 proteins are colored in magenta and blue for 2019-nCoV and in green and red for SARS-CoV, respectively. **(B)** The binding site residues on the RBD that are within 5.0 Å form the ACE2, where the hydrophobic residues on the 2019-nCoV RBD are highlighted in red. **(C)** Part of the sequence alignment for the spike proteins between 2019-nCoV and SARS-CoV, where the RBD residues are highlighted in yellow and the binding site residues are highlighted in magenta, respectively.

### 3.2 The RBD-ACE2 complex: 2019-nCoV binds ACE2 with higher affinity than SARS-CoV

We have run a long-time MD simulation to generate the trajectories of the RBD-ACE2 complex system for 2019-nCoV and SARS-CoV. The binding free energies were calculated using the MM-GBSA model by the “MMPBSA.py” script in the AMBER package. Table 1 shows a comparison of the RBD-ACE2 binding free energies for 2019-nCoV and SARS-CoV. It can be seen from the table that the binding free energy of the RBD-ACE2 interaction for 2019-nCoV is −50.43 kcal/mol, which is significantly lower than the binding free energy of the RBD-ACE2 interaction for SARS-CoV (−36.75 kcal/mol). In other words, 2019-nCoV binds human ACE2 with a significantly higher affinity than SARS-CoV. This result is consistent with the current fact that 2019-CoV is much more infectious than SARS-CoV. Very recently, experimental studies also showed that 2019-CoV could bind human ACE2 with a higher affinity than SARS-CoV [22]. Further examination of the binding free energy contributions reveals that the higher binding affinity of 2019-nCoV than SARS-CoV is mostly attributed to the solvation energy contribution Δ*G*_solv_ (674.97 vs. 696.56 kcal/mol), whereas 2019-nCoV has a higher binding free energy in vacuum Δ*G*_gas_ than SARS-CoV (−725.41 vs. −733.31). In other words, 2019-nCoV tends to bind human ACE2 better than SARS-CoV in the water, while SARS-CoV would bind to human ACE2 better than 2019-nCoV in the gas. Further investigation is needed to confirm such binding differences.

**Table 1:** The binding free energies calculated from MD simulations for the RBD-ACE2 interactions of 2019-nCoV and SARS-CoV, where Δ*G*_gas_ is the interaction energy change in the gas and Δ*G*_solv_ is the solvation energy change in the solvent upon binding, respectively. All units are reported in kcal/mol.

|  | 2019-nCoV |  |  | SARS-CoV |  |  |
| --- | --- | --- | --- | --- | --- | --- |
|  | Average | Std. Dev. | Std. Err. of Mean | Average | Std. Dev. | Std. Err. of Mean |
| $\Delta G_{\text{gas}}$ | -725.4066 | 31.2987 | 1.3997 | -733.3113 | 33.8176 | 1.5124 |
| $\Delta G_{\text{solv}}$ | 674.9721 | 29.5988 | 1.3237 | 696.5572 | 29.3955 | 1.3146 |
| $\Delta G_{\text{total}}$ | -50.4345 | 5.5811 | 0.2496 | -36.7541 | 7.4944 | 0.3352 |

### 3.3 The spike protein: 2019-nCoV is more stable than SARS-CoV

The spike protein on the coronavirus envelope is a trimeric protein. This protein is critical for the vitality of coronaviruses because it is not only an important component for the virus particle but also plays a crucial role in attaching host cells and fusing the membranes [21]. In addition, the spike protein also determines the solubility of coronavirus particles and thus the viral infectivity because the spike protein is the largest protein located on the membrane surface. Therefore, the spike protein is directly related to the stability and functionality of coronaviuses. Here, we have run a lengthy MD simulation to study the trimeric spike proteins of 2019-nCoV and SARS-CoV.

Table 2 gives a comparison between the free energies of the spike proteins for 2019-nCoV and SARS-CoV. It can be seen from the table that the spike protein of 2019-nCoV has a significantly lower total free energy (*G*_total_ = −67303.28 kcal/mol) than the spike protein of SARS-CoV (*G*_total_ = −63139.96 kcal/mol) (Table 2). These results suggest that 2019-nCoV is more stable and can survive a significantly higher temperature than SARS-CoV. This may also partly explain the higher infectivity of 2019-nCoV than SARS-CoV because 2019-nCoV would have a higher virus vitality than SARS-CoV in the same temperature.

**Table 2:** The free energies calculated from MD simulations for the spike proteins of 2019-nCoV and SARS-CoV, where *G*_gas_ is the interaction energy in the gas and *G*_solv_ is the solvation energy in the solvent, respectively. All units are reported in kcal/mol.

|  | 2019-nCoV |  |  | SARS-CoV |  |  |
| --- | --- | --- | --- | --- | --- | --- |
|  | Average | Std. Dev. | Std. Err. of Mean | Average | Std. Dev. | Std. Err. of Mean |
| $G_{\text{gas}}$ | -36405.4424 | 294.4540 | 29.4454 | -32053.4298 | 270.8693 | 27.0869 |
| $G_{\text{solv}}$ | -30897.8370 | 283.0603 | 28.3060 | -31086.5339 | 236.9148 | 23.6915 |
| $G_{\text{total}}$ | -67303.2794 | 171.9868 | 17.1987 | -63139.9637 | 199.4728 | 19.9473 |

The lower free energy of the 2019-nCoV spike protein may result from the virus evolution of SARS-CoV because SARS-like coronaviruses normally originate from bats that are known to have a higher body temperature than humans. In other words, 2019-nCoV and other SARS-like coronavirus would have evolved to achieve a lower free energy for their spike proteins by recombination or mutations so that they can survive in high-temperature animals like bats [1]. In addition, the free energy decomposition also shows that the lower free energy of 2019-nCoV spike protein than SARS-CoV spike protein is mainly attributed to the free energy in vacumm *G*_gas_ (−36405.44 vs. −32053.43 kcal/mol), whereas their solvation energies *G*_solv_ are comparable (−30897.84 vs. −31086.53 kcal/mol) (Table 2). This may reflect an evolution trend for SARS-like coronaviruses, i.e. favoring the internal interactions between residues instead of the solvation energy. This kind of evolution would be also beneficial because such kinds of coronaviruses would be more robust and able to survive in both the air and solvent.

### 3.4 The RBD domain: 2019-nCoV is more temperature-sensitive than SARS-CoV

Coronaviruses uses the spike protein to attach host cells by binding the host cell receptor. Therefore, the receptor binding domain (RBD) of the spike protein is critical for coronaviruses to infect host cells. Here, we have run lengthy MD simulations to investigate the structural properties of the RBD proteins for 2019-nCoV and SARS-CoV. Table 3 shows a comparison between the free energies of the RBD proteins for 2019-nCoV and SARS-CoV. Similar to the findings in the spike protein as detailed in the last section (Table 2), the RBD protein of 2019-nCoV also shows a significantly lower free energy than that of SARS-CoV (−4090.04 vs. −3617.73 kcal/mol) (Table 3), which may also be understood by the evolution pressure from the high-temperature host environment. However, unlike in the spike protein where the free energy difference between 2019-nCoV and SARS-CoV is mostly attributed the inter-residue interactions in vacuum (*G*_gas_), here in the RBD protein, the free energy difference between 2019-nCoV and SARS-CoV comes from both the free energy in vacuum *G*_gas_ (−2104.37 vs. −1703.66 kcal/mol) and solvation energy *G*_solv_ (−1985.68 vs. −1914.07 kcal/mol) (Table 3). The lower solvation energy of 2019-nCoV than SARS-CoV in the RBD may be understood because the RBD must move up away from the spike protein and into the water in order to bind human ACE2 [20]. In other words, 2019-nCoV would have evolved to be more soluble so that it can move up and bind the ACE2 more easily. The better solubility of the RBD of 2019-nCoV than SARS-CoV may also contribute to part of the higher infectivity of 2019-nCoV than SARS-CoV.

**Table 3:**
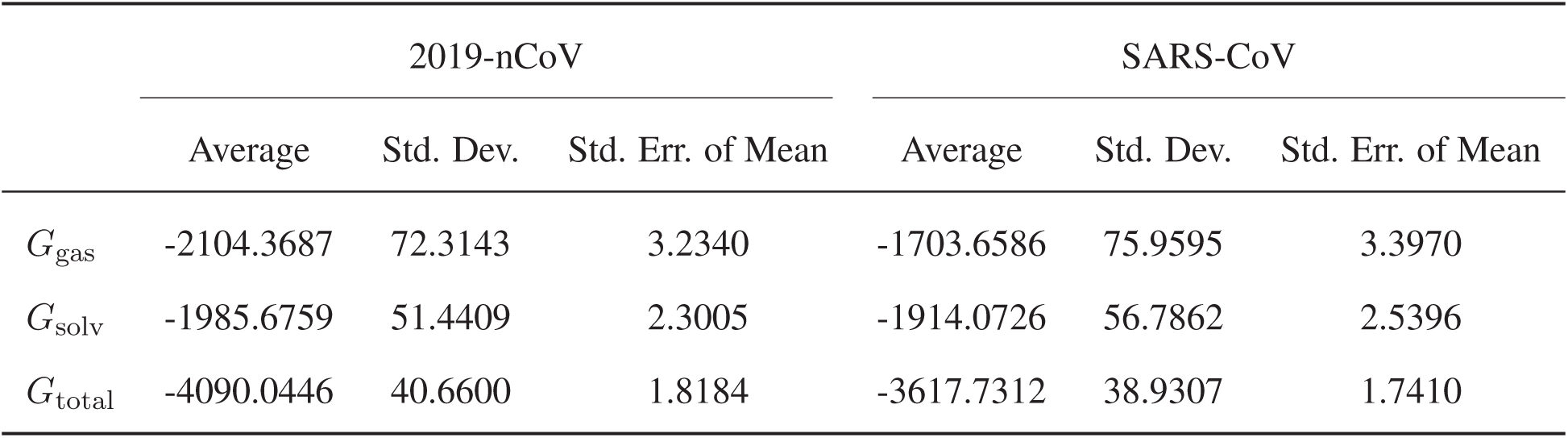
The free energies calculated from MD simulations for the RBD proteins of 2019-nCoV and SARS-CoV, where *G*_gas_ is the interaction energy in the gas and *G*_solv_ is the solvation energy in the solvent, respectively. All units are reported in kcal/mol.

Protein flexibility is a critical factor in binding as it may not only change the binding interface between two proteins but also be an important contribution to the entropy penalty upon binding. Therefore, we have investigated the protein flexibility of the RBD domains for 2019-nCoV and SARS-CoV by analyzing their coordinate trajectories. Figure 2 show two ensembles of selected trajectories over a period of 10ns simulations for the RBD domains of 2019-nCoV and SARS-CoV. The figure also gives a comparison of the root mean square fluctuations (RMSF), a rough measurement of protein flexibility, for 2019-nCoV and SARS-CoV. It can be seen from the figure that the RBD of 2019-nCoV shows a significantly higher RMSF than that of SARS-CoV. In other words, the RBD of 2019-nCoV is much more flexible than the RBD of 2019-nCoV. The flexibility is especially higher near the binding site than other regions (Figure 2A). That means that 2019-nCoV must overcome much more entropy penalty than SARS-CoV when binding to human ACE2. As we know, the binding free energy between two proteins can be expressed as, Δ*G* = Δ*E* − *T* Δ*S*, where Δ*E* is the interaction energy, Δ*S* is the entropy, and *T* is the temperature of the system. As Δ*S* is negative, the binding free energy will become higher and the binding will become weaker with the increasing temperature. Therefore, the RBD-ACE2 binding affinity for 2019-nCoV is expected to decrease much faster than that for SARS-CoV when the temperature increases. In other words, 2019-nCoV is much more temperature-sensitive than SARS-CoV. Namely, 2019-nCoV will decrease its infection ability much faster than SARS-CoV when the temperature rises. Therefore, it is expected that 2019-nCoV will become less infectious compared to SARS-CoV, and the disease prevention and control for 2019-nCoV will get easier when the weather gets warmer/hotter, although the drug and vaccine development targeting the RBD protein will be more challenging because of the protein flexibility at the binding site.

**Figure 2:**
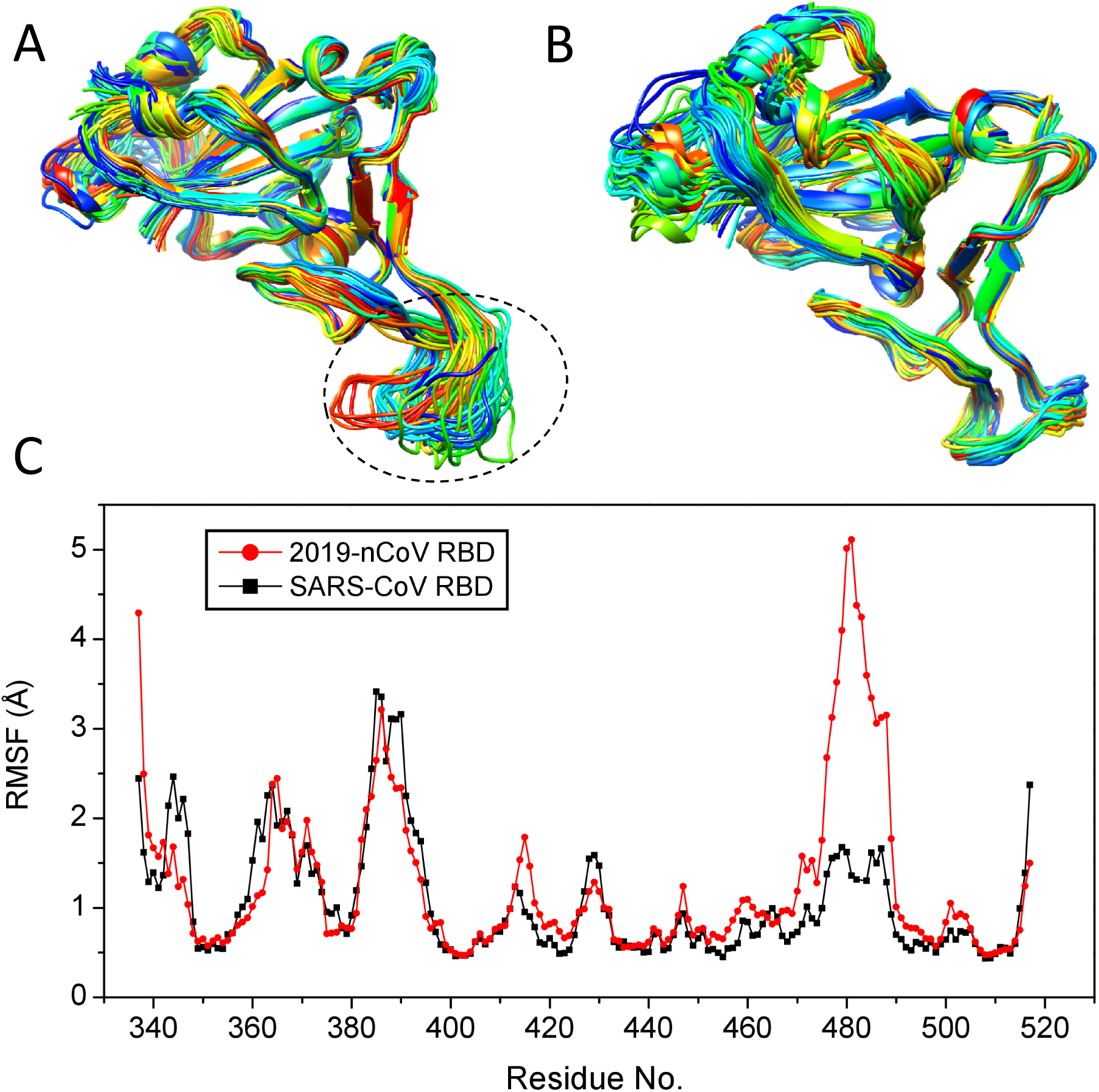
The conformational flexibility of the RBD protein. **(A)** The 50 representative trajectories of the 2019-nCoV RBD protein over a 10ns MD simulation, where the highly flexible region (residues 470-490) is indicated by a circle. **(B)** The 50 representative trajectories of the SARS-CoV RBD protein over a 10ns MD simulation. **(C)** The RMSFs of the 2019-nCoV and SARS-CoV RBD proteins, where the residue numbering is taken from the 2019-nCoV RBD and the data of SARS-CoV are then aligned to those of 2019-nCoV according to the sequence alignment.

## 4 CONCLUSIONS

In this study, we have extensively studied the RBD-ACE2 complex, spike protein, and free RBD protein systems of 2019-nCoV and SARS-CoV through protein-protein docking and MD simulations. It was found that 2019-nCoV has a higher binding affinity with human ACE2 than SARS-CoV, explaining the fact that 2019-CoV is much more infectious than SARS-CoV. The spike protein of 2019-nCoV also shows a lower free energy than that of SARS-CoV, suggesting that 2019-nCoV is more stable and may be able to survive a higher temperature than SARS-CoV. This may also explain the bat origin of 2019-nCoV, as bats have a higher body-temperature than humans. In addition, the RBD of 2019-nCoV exhibits a significantly higher flexibility than that of SARS-CoV, especially near the binding site. That indicates that 2019-nCoV must overcome a higher entropy penalty in order to bind ACE2 and is thus more temperature-sensitive than SARS-CoV. Therefore, with the rising temperature, 2019-nCoV is expected to decrease the infection ability much faster and become much less infectious than SARS-CoV, which would make the disease prevention and control of 2019-nCoV easier. Taking the above results together, unlike SARS-CoV that is gone after 2003, 2019-nCoV might survive the high-temperature environment like Summer in which the virus is not active/infectious due to the high flexibility in the RBD, and then become infectious when the temperature is low in the Winter. These findings will have a far-reaching implication for disease prevention and control as well as drug and vaccine development for 2019-nCoV.

## ACKNOWLEDGEMENTS

This work is supported by the National Natural Science Foundation of China (grant Nos. 31670724) and the startup grant of Huazhong University of Science and Technology.

